# Transcriptional heterogeneity of *Cryptococcus gattii* VGII compared with non-VGII lineages underpins key pathogenicity pathways

**DOI:** 10.1101/396796

**Authors:** Rhys A. Farrer, Christopher B. Ford, Johanna Rhodes, Toni Delorey, Robin C. May, Matthew C. Fisher, Elaine Cloutman-Green, Francois Balloux, Christina A. Cuomo

## Abstract

*Cryptococcus gattii* is a pathogenic yeast of humans and other animals, which causes disease predominantly in immunocompetent hosts. Infection begins when aerosolized yeast or spores enter the body, triggering an immune response, including engulfment by macrophages. To understand the early transcriptional signals in both the yeast and its mammalian host, we performed a time-course dual RNA-seq experiment for four lineages of *C. gattii* (VGI-IV) interacting with mouse macrophages at 1hr, 3hr and 6hr post infection. Comparison of *in vitro* to *ex vivo* gene expression indicates lineage VGII is transcriptionally divergent to non-VGII lineages, including differential expression of genes involved in capsule synthesis, capsule attachment and ergosterol production. Various paralogs demonstrate sub-functionalisation between lineages including an upregulation of capsule biosynthesis-related gene *CAP2*, and downregulation of *CAP1* in VGIII. Isolates also compensate for lineage-specific gene-losses by over-expression of genetically similar paralogs, including an over-expression of capsule gene *CAS3* in VGIV having lost *CAS31*. Differential expression of one in five *C. gattii* genes was detected following co-incubation with mouse macrophages; all isolates showed high induction of oxidative-reduction functions and a downregulation of capsule attachment genes. We also show that VGII switches expression of two laccase paralogs (from *LAC1* to *LAC2*) during co-incubation of macrophages. Finally, we found that mouse macrophages respond to all four lineages of *C. gattii* by upregulating FosB/Jun/Egr1 regulatory proteins at early time points. This study highlights the evolutionary breadth of expression profiles amongst the lineages of *C. gattii* and the diversity of transcriptional responses at this host-pathogen interface.

**Importance:** The transcriptional profiles of related pathogens and their response to host induced stresses underpin their pathogenicity. Expression differences between related pathogens during host interaction can indicate when and how these genes contribute to virulence, ultimately informing new and improved treatment strategies for those diseases. In this paper, we compare the transcriptional profiles of five isolates representing four lineages of *C. gattii* in rich media. Our analyses identified key processes including cell capsule, ergosterol production and melanin that are differentially expressed between lineages, and we find that VGII has the most distinct profile in terms of numbers of differentially expressed genes. All lineages have also undergone sub-functionalisation for various paralogs including capsule biosynthesis and attachment genes. Most genes appeared down-regulated during co-incubation with macrophages, with the largest decrease observed for capsule attachment genes, which appears coordinated with a stress response, as all lineages also upregulated oxidative stress response genes. Furthermore, VGII upregulated many genes that are linked to ergosterol biosynthesis and switched expression of the laccase *LAC1* to *LAC2 ex vivo*. Finally, we saw a pronounced increase in the FosB/Jun/Egr1 regulatory proteins at early time points in bone marrow derived macrophages, marking a role in the host response to *C. gattii*. This work highlights the dynamic roles of key *C. gattii* virulence genes in response to macrophages.

## Introduction

Infectious diseases impose a huge burden on human society. In recent years, fungi have gained widespread attention for their ability to threaten both animal and plant species across a global scale (1, 2). However, many features of the fungal genome and transcriptome that enable the infection of diverse hosts and ecological niches remain largely unexplored, especially for emerging pathogens (3, 4). One such example is the basidiomycete yeast *Cryptococcus gattii*, which can cause pneumonia and meningoencephalitis in humans (5). While *C. gattii* causes less overall global morbidity than its sibling species *C. neoformans*, lineages have emerged with hypervirulent clinical phenotypes, a process that was most strikingly observed in the Pacific Northwest USA in the late 1990’s (6). *C. gattii* is comprised of four genetically distinct lineages designated as VGI to VGIV molecular types, of which lineage VGII is associated with the highly virulent subtypes seen in the USA. These lineages are sufficiently divergent to include 737 lineage-specific gains and/or losses of genes across all four lineages (approximately 4% of the genes in any given isolate), including DNA transposons, genes involved in the response to oxidative stress and import into mitochondrial inner membrane (7).

In addition to genetic differences between lineages, changes in gene regulation underlie important morphological and physiological traits in fungal pathogens (8, 9). However, the mechanisms of gene-regulation, evolution of gene networks, and rewiring of transcriptional modules within lineages remain largely uncharacterized for many infectious diseases including *C. gattii.* New virulent lineages can emerge through mutation and/or recombination events (10), which in turn lead to different transcriptional profiles and enhanced virulence profiles, such as the hyper-virulent *C. gattii* VGIIa sub-lineage in the Pacific Northwest (11), which descends from two alpha mating-type parents (6) and is characterized by enhanced intracellular parasitism (11).

Previous studies have demonstrated unique expression profiles among different isolates of *Cryptococcus* within varying conditions. For example, in *C. neoformans* reference isolate H99, genes encoding membrane transporters for nutrients, general metabolism, and oxidative stress response have been shown to be upregulated in the presence of macrophages (12) and amoeba (13). Janbon *et al.* also characterized differential expression of transporters, transcription factors and genes involved in lipid metabolism in *C. neoformans* between rich media, limited/starved media and pigeon guano (14). In addition to analysing protein coding gene expression, Janbon *et al.* also identified nearly 1,200 miscellaneous RNAs that may perform a range of functions including morphogenesis *via* small ORFs, or ncRNA with structural or regulatory roles. Concurrently, Chen *et al.* described an increase in genes involved in metabolic processes, alkaline response, salt tolerance and oxidative stress for two *C. neoformans* isolates in cerebral spinal fluid compared to growth in rich media (15).

Few studies have focused on gene expression in *C. gattii*, although notably Ngamskulrungroj *et al.* identified an upregulation of laccase genes involved in melanin formation, and other genes involved in cell wall assembly and metabolism in *C. gattii* VGIIa outbreak strain R265, relative to the non-outbreak *C. gattii* VGIIb strain in minimal medium (16). In this study, we extend the description of *C. gattii* gene expression levels to characterise variation across all four major lineages of *C. gattii*, including the outbreak strain VGIIa R265 and a non-outbreak isolate ENV152 also from the VGIIa subgroup, both *in vitro*, and at three early time points (1hr, 3hr and 6hr) following co-incubation and engulfment by murine bone marrow derived macrophages (BMDMs). Furthermore, we describe concurrent expression changes in host macrophages during *C. gattii* co-incubation, demonstrating the potential for the simultaneous profiling of both host and pathogen responses.

## Results

To identify mammalian and *Cryptococcus gattii* genes that are activated during infection, we performed RNA-seq across five *C. gattii* isolates representing four lineages (VGI, VGII, VGIII and VGIV), including an environmental and clinical isolate from VGII, both *in vitro* and at time points 1, 3 and 6 hours post co-incubation with BMDMs (*ex vivo*). Each of the sequencing runs (*n*=40) yielded between 904,770 and 17.6 million 30mer reads (10.9 Gigabase of total sequence) (**Table S1, Fig. S1, Fig. S2, Fig. S3**), except for VGIV at *t*6, which failed to yield sequencing data. Most of the sequences derived from mouse (84-99% for each dataset), while *C. gattii* reads ranged from up to 56.97% (*in vitro* without macrophages) to between 0.02-1.44% reads *ex vivo* (mean of 38,949 reads) (**Fig. S2**). The *C. gattii* datasets formed two main clusters: 1) *t*0 (*in vitro*) for all isolates, probably owing to the huge difference in read counts between samples and 2) VGII isolates at any time point (**Fig. 1a**). Conversely, gene expression in mouse macrophages clustered by time point (**Fig. 1b**).

**Figure 1.**
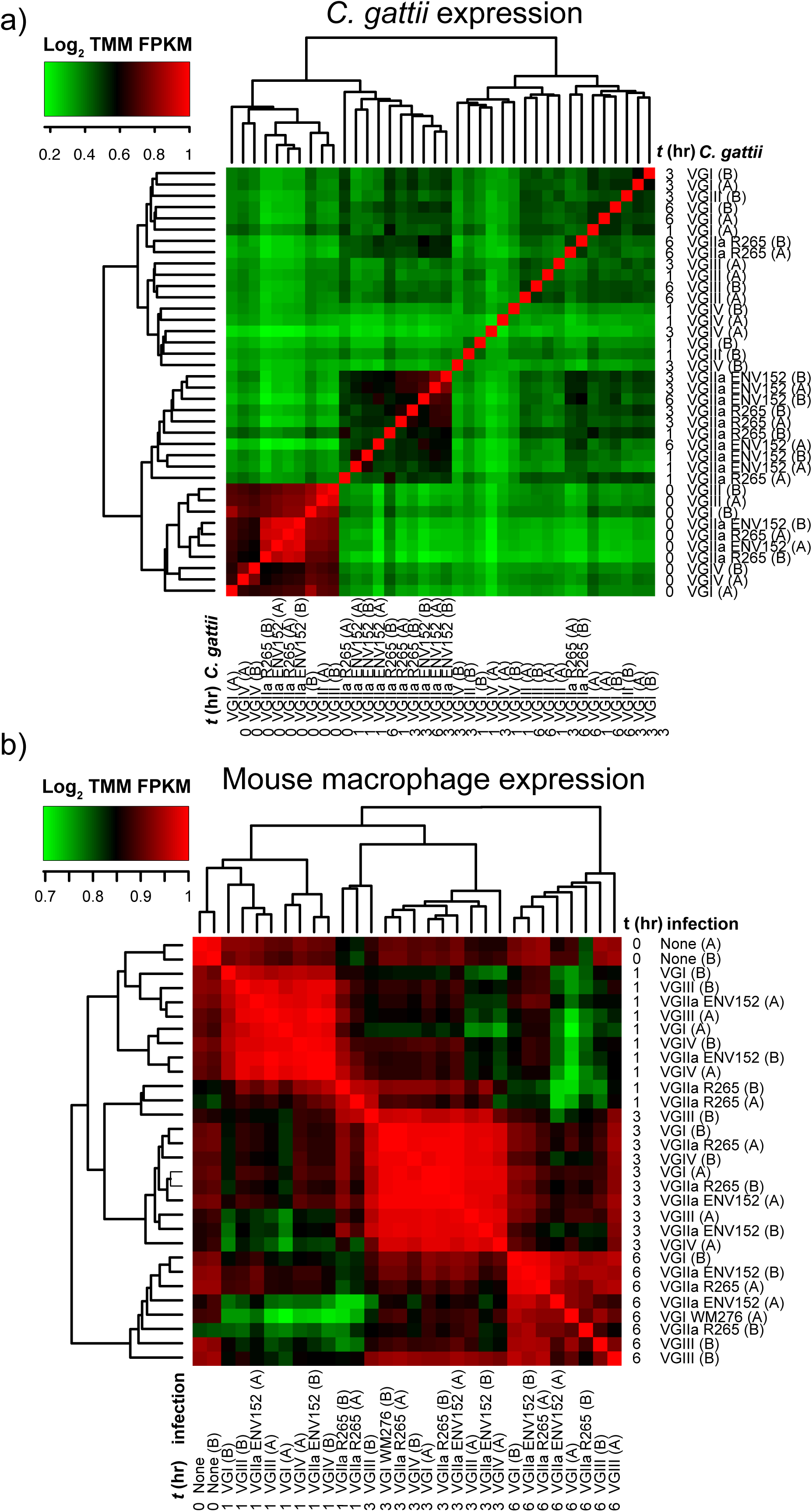
Heat maps of the Log_2_ fold change in the trimmed mean of M-values (TMM) normalized Fragments Per Kilobase of transcript per Million mapped reads (FPKM) of *C. gattii* (a) and Mouse BMDM (b) transcripts per sample.

Differential expression was computed using the quantile-adjusted conditional maximum likelihood (qCML) method implemented in EdgeR (17), requiring a false discovery rate (FDR) *p*-value <1e^-3^ and at least fourfold change in Trimmed Mean of M-values (TMM) normalized Fragments Per Kilobase of transcript per Million Mapped reads (FPKM), to be considered as a significantly differentially expressed gene (DEG). To check for the effect of different read-depths on DEG prediction, subsets were created and re-analysed. This analysis identified consistent numbers of DEGs *in vitro*, and more variable numbers between infection time-points (**Fig. S4).**

### VGII is transcriptionally divergent *in vitro* from other *C. gattii* lineages

Pairwise expression values for all *C. gattii* isolates *in vitro* (*t*0) revealed 524 DEGs between 1 or more isolates, indicating that nearly one in ten (8.1%) of all *C. gattii* protein-coding genes situated throughout the genome (*n*=6,456) are uniquely differentially regulated among the four divergent lineages, despite being all cultured in the same rich media at 37°C with 5% CO_2_ (**Fig. 2a, Fig. 2e, Sup. Dataset 1**). A *G*-test of goodness-of-fit based on just the numbers of *in vitro* DEGs per lineage (not considering specific genes) suggested differences (*G* = 328.52, X-squared df = 4, *p*-value < 2.2e^-16^), and pairwise comparisons using G-tests with Bonferroni multiple correction also showed differences between VGII isolates compared with VGI, VGII ENV152 and VGIV, but not VGII R265 (*p*-value < 2.2e^-16^). VGIV compared with VGI or VGIII were also highly distinct for numbers of DEGs (*p*-value < 2.2e^-16^).

Of the 524 *in vitro* DEGs, 203 were differentially regulated in both VGII isolates, 245 additional genes were differentially regulated for only an individual VGII isolate, and 50 were differentially regulated inconsistently across VGII (e.g. upregulated in VGIV *vs* VGII R265 and upregulated in VGI *vs* both VGII isolates). While some of these genes were also differentially expressed in other lineages, only 5% (*n*=27) of *in vitro* DEGs were unique to VGI, VGIII or VGIV isolates, suggesting VGII harbors distinct expression profiles compared with the other lineages. Furthermore, the greater number of genes uniquely differentially expressed within isolates of the VGII isolates suggests that substantial differences exist even within the VGIIa sub-lineage, possibly owing to genetic variation generated from their different sources of origin (environmental or clinical). Furthermore, VGII isolates were distinct by Principle Component Analysis (PCA) (**Fig. 2c**), while VGI and VGIV were not.

**Figure 2.**
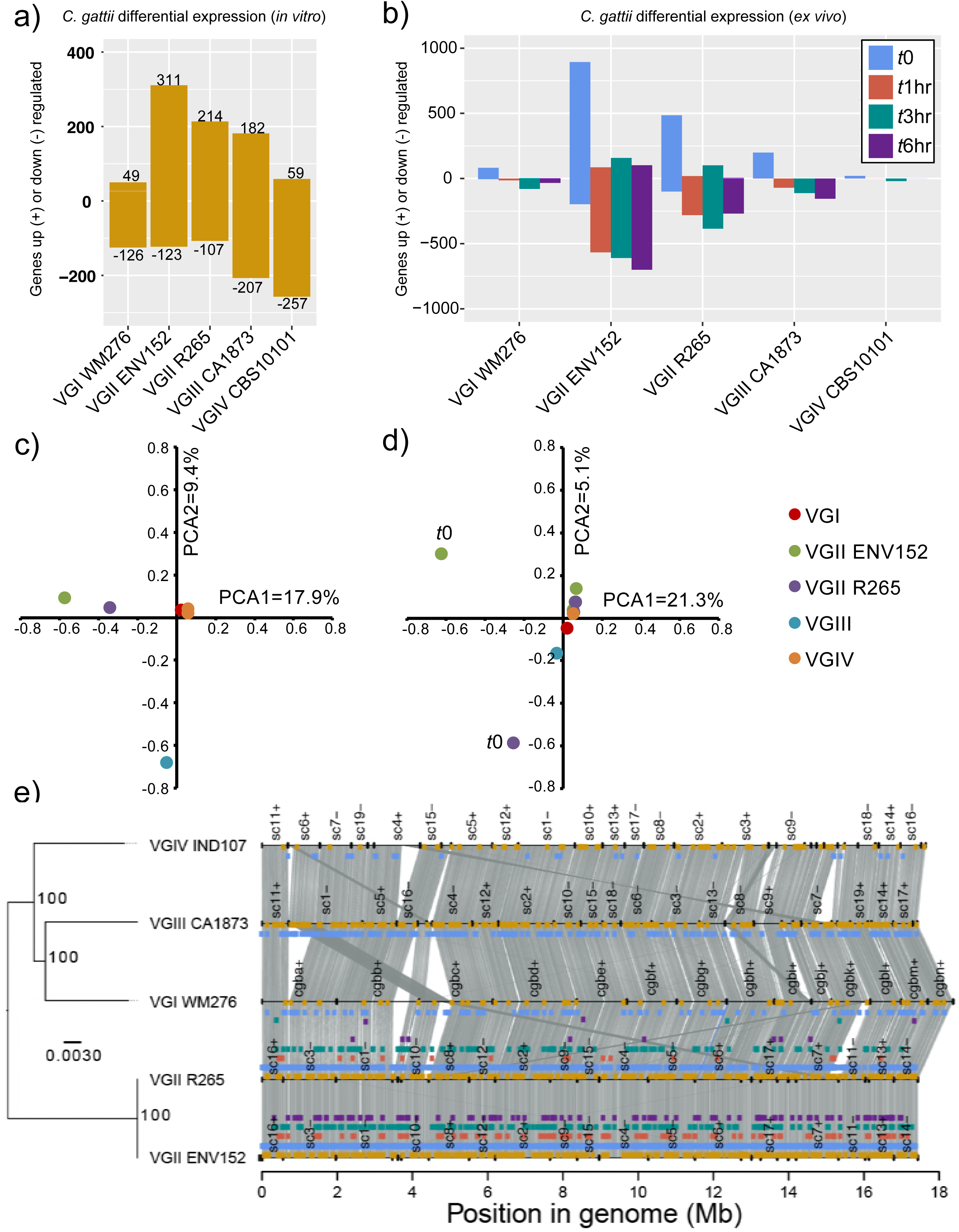
The number of *Cryptococcus gattii* genes up/downregulated between lineages *in vitro* (a), between time-points (b), Principle Component Analysis (PCA) of *in vitro* (c) and between time-point (d) differentially expressed genes and the locations of those genes in their genomes (synteny plotted using Synima (39)) alongside a phylogenetic tree constructed with RAxML (GTRCAT) with 1000 bootstrap support (shown as asterisks) (e). Genes are considered differentially expressed where FDR *p* value<0.001 and >fourfold TMM FPKM.

To understand the role that differential expression may have on each isolate *in vitro*, we opted for both a targeted approach (looking at known genes of interest including 35 capsule biosynthesis genes, 40 capsule attachment and cell-wall remodelling genes and 20 ergosterol genes (18) based on their orthology to *C. neoformans* H99 (7)) and a non-targeted approach (GO-term and PFAM enrichment). Of the 524 unique DEGs, 8/35 were involved in capsule biosynthesis, 10/40 in capsule attachment and cell-wall remodelling, 4/20 in ergosterol production, and both laccase genes were differentially expressed in at least one pairwise comparison (**Fig. 3**). These gene categories are therefore enriched for DEGs based on a Hypergeometic Test (P[X > x]) = *P*=1.207e^-07^).

**Figure 3.**
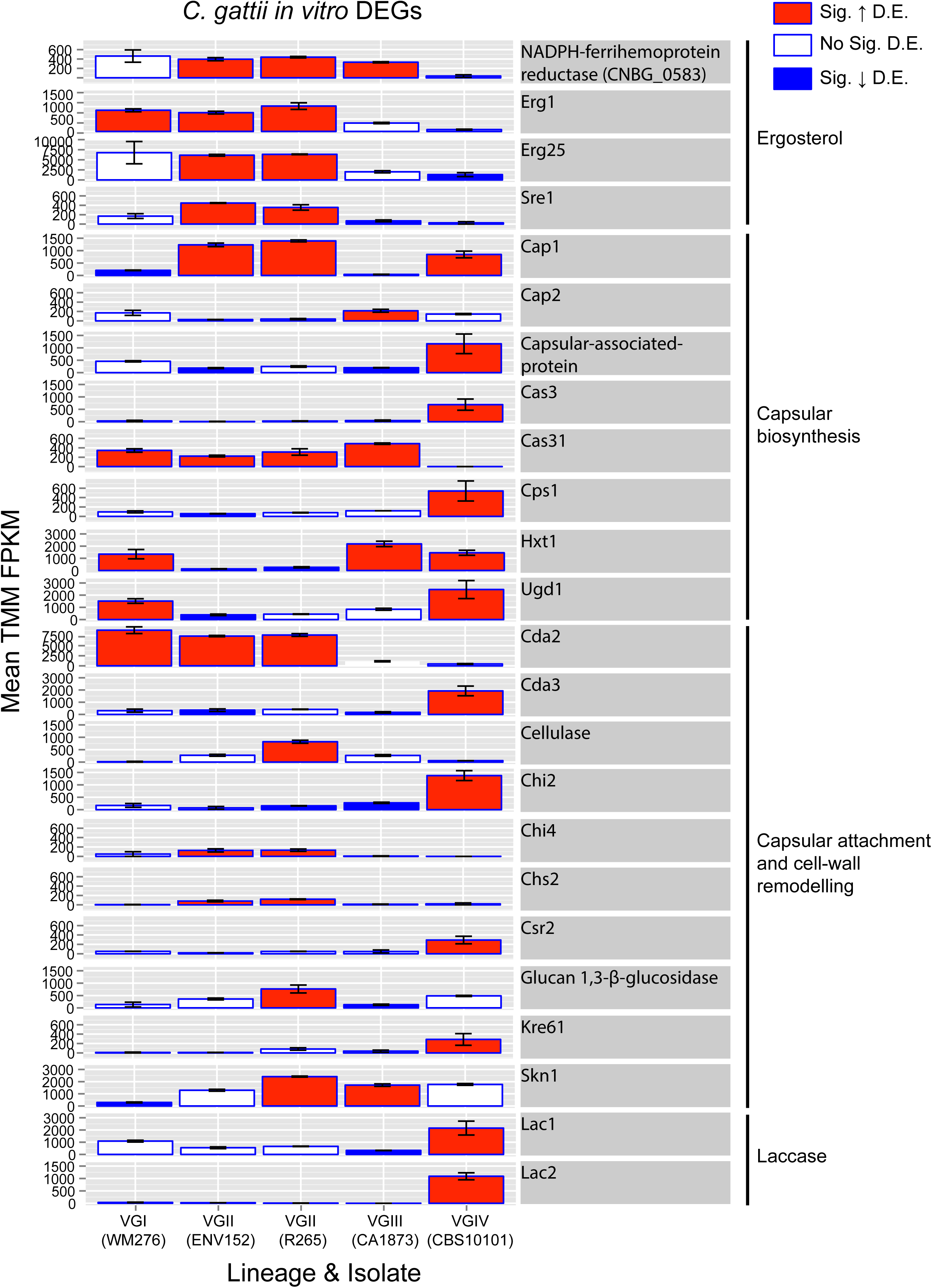
Bar charts showing mean expression (TMM FPKM) of ergosterol, capsular biosynthesis, capsular attachment and laccase genes between the five isolates of *Cryptococcus gattii in vitro*. Red=Sig. upregulated, Blue=Sig. downregulated. Genes are considered differentially expressed where FDR *p* value<0.001 and >fourfold TMM FPKM. Error bars show the range of TMM FPKM values between the two replicates.

### *In vitro C. gattii* lineages have distinct expression for capsule biosynthesis and attachment genes

Capsule biosynthesis genes were differentially expressed between lineages *in vitro* (**Fig. 3**), including *CAS3* which was upregulated in VGIV compared with all the other isolates. *CAS3* mutants have a reduced capsule when combined with *cas31*Δ, *cas32*Δ, or *cas33*Δ mutants, and have a partial defect in O-acetylation leading to reduced overall levels of this modification (18, 19). The closely related paralog *CAS31* is absent in VGIV (but present in the other three lineages) (**Fig. S5**), and as such has no detectable expression (zero TMM FPKM for both replicates) – manifesting as downregulated compared with each of the other lineages. It is therefore possible that VGIV is over-expressing *CAS3* to compensate for its *CAS31* deletion or disruption and demonstrating sub-functionalisation of these paralogs. Furthermore, *cas31*Δ have previously been shown to manifest minor differences in GXM composition in *C. neoformans* (18, 19), which may also manifest in *C. gattii* VGIV. Similarly, *CAP1* is upregulated in VGII and VGIV isolates relative to VGI and VGIII isolates, while the close paralog *CAP2* is upregulated by VGIII compared with VGII. Other differentially expressed capsule biosynthesis genes may not be fully compensated by genetically similar paralogs, such as the hexose transporter *HXT1* downregulated by both VGII isolates.

Capsule attachment and cell-wall remodelling genes were also differentially expressed between lineages of *C. gattii* (**Fig. 3**). For example, chitin synthases *CHI4*, and *CHS2* were upregulated in both VGII isolates. Chitinase *CHI2* was upregulated in VGIV. Chitin deacetylase 1 (*CDA1*) was highly expressed by all isolates *in vitro* (TMM FPKM ranging from 2265-932 across all replicates) and was not identified as differentially expressed. However, *CDA2* was upregulated in VGI and VGII isolates compared with VGIII and VGIV. Meanwhile, *CDA3* was upregulated in VGIV compared with VGII and VGIII (perhaps again showing a change in paralog expression regulation). Both *CDA2* and *CDA3* mutants have increased capsule size when combined with *cda1*Δ (18), however, *CDA1* was not found to be differentially expressed.

### Differential expression of known drug targets and virulence genes

In common with most fungi, ergosterol is found in the membrane of *C. gattii* and is a key target for numerous anti-fungal drugs including fluconazole and amphotericin B (20). The VGII lineage had significantly higher expression for several genes involved in ergosterol production including NADPH-ferrihemoprotein reductase, *ERG1* (squalene monooxygenase), *ERG25* (methylsterol monooxygenase), and *SRE1*. Each of these genes are linked to drug-resistance or the oxidative stress response, for example *ERG1* mutants have increased fluconazole susceptibility in *Candida glabrata* (21), *ERG25* has a moderate susceptibility to hypoxia and the E.R. stress-inducing agent DTT in *Aspergillus fumigatus* (22, 23), and *SRE1* is required for hypoxic induction of genes encoding for oxygen-dependent enzymes involved in ergosterol synthesis in *C. neoformans* (24). The biological significance of these expression differences are unclear, but could manifest in lineage-specific drug resistance variation.

Laccases are cell wall enzymes that catalyse melanin to protect *Cryptococcus* from various stresses including oxidative stress, and are therefore considered important virulence factors (25). Laccase production in both *C. gattii* and *C. neoformans* is controlled by two cell wall enzymes that control melanin production (25). VGIV significantly upregulates both copies (*LAC1* and *LAC2*) *in vitro* compared with either VGIII or all the other isolates respectively. While *LAC1* and *LAC2* are flanking each other in *C. neoformans*, *C. gattii* has an additional gene CNBG_2145 encoding a hypothetical protein with no functional annotation (PFAM, GO, KEGG) in the middle. This gene, unlike *LAC1* and *LAC2*, was not consistently expressed by any of the lineages in any condition.

Differential expression among isolates *in vitro* were also enriched (Two-tailed Fisher’s exact test with *q*-value FDR) for several GO-terms compared with the remaining genes (**Table 1**). Enriched terms included oxidative reduction (*q*=1.85E^-07^) and oxidoreductase activity (*q*=0.011). Genes with the oxidative reduction GO-term were both up and down regulated by each isolate, including ferric reductases, metalloreductases, nitric oxide dioxygenases, acidic laccases, oxidoreductases and various dehydrogenases. VGII isolates had the greatest number of up-regulated genes assigned an oxidative reduction function: VGII ENV152 (*n*=39), VGII R265 (*n*=23), compared with VGIII (*n*=18), VGIV (*n*=11) and VGI (*n*=7). All R265 genes were also found in ENV152 apart from the single gene CNBG_2804 (hypothetical protein with a DUF455 domain). The remaining 17 uniquely upregulated genes in ENV152 included five dehydrogenases (methylmalonate-semialdehyde, 3-hydroxyacyl-CoA, glutaryl-CoA, glyceraldehyde-3-phosphate, glutamate), Fe-Mn family superoxide dismutase and the ferric reductase transmembrane component 4. Only a single PFAM, PF00083.19 sugar (and other molecule) transporter, was enriched from the *in vitro* comparisons. The ability of *C. gattii* to respond to host and environment derived oxidative stress is well described (22, 26, 27) and it is noteworthy that differences in expression level were found between isolates and lineages even in *in vitro* conditions.

**Table 1.** Enrichment of functional annotation for in vitro and ex vivo comparisons was determined by a two-tailed Fisher exact test with *q*-value (Storey-Tibshirani) FDR, where count 1 are number of terms and parent terms in the differentially expressed set, count 2 are the remaining terms and parent terms in the non-differentially expressed set. The uncorrected fisher *p*-values, corrected *q*-values, relative proportion and description are provided for each term (requiring *q*-values of <0.05 for each).

| Sig. enriched terms from *in vitro* comparisons |  |  |  |  |  |  |
| --- | --- | --- | --- | --- | --- | --- |
| GO/PFAM | count 1 | count 2 | fisher *p* | *q* value | rel. prop | GO description |
| GO:0055114 | 64 | 256 | 1.13E-10 | 1.85E-07 | 2.54 | oxidation reduction |
| GO:0016638 | 5 | 2 | 1.01E-04 | 1.11E-02 | 25.41 | oxidoreductase activity, acting on the CH-NH2 group of donors |
| GO:0050660 | 13 | 32 | 9.79E-05 | 1.11E-02 | 4.13 | FAD binding |
| PF00083.19 | 18 | 54 | 2.26E-05 | 2.66E-02 | 3.56 | Sugar_tr [Sugar (and other) transporter] |
| GO:0004553 | 12 | 33 | 4.15E-04 | 3.25E-02 | 3.7 | hydrolase activity, hydrolyzing O-glycosyl compounds |
| GO:0005975 | 25 | 116 | 7.41E-04 | 4.86E-02 | 2.19 | carbohydrate metabolic process |

Sig. depleted terms from *in vitro* comparisons
| GO term | count 1 | count 2 | fisher *p* | *q* value | rel. prop | GO description |
| --- | --- | --- | --- | --- | --- | --- |
| GO:0043234 | 3 | 211 | 3.35E-06 | 6.14E-04 | 0.14 | protein complex |
| GO:0034645 | 11 | 316 | 6.33E-05 | 9.42E-03 | 0.35 | cellular macromolecule biosynthetic process |
| GO:0006396 | 2 | 142 | 2.60E-04 | 2.25E-02 | 0.14 | RNA processing |
| GO:0003723 | 2 | 137 | 3.78E-04 | 3.11E-02 | 0.15 | RNA binding |
| GO:0005737 | 35 | 597 | 7.23E-04 | 4.86E-02 | 0.6 | cytoplasm |
| GO:0015031 | 2 | 126 | 7.96E-04 | 4.86E-02 | 0.16 | protein transport |

Sig. enriched terms from *ex vivo* comparisons
| GO/PFAM | Isolate | count 1 | count 2 | *q* value | rel. prop | GO description |
| --- | --- | --- | --- | --- | --- | --- |
| GO:0055114 | VGI WM276 | 18 | 302 | 3.56E-05 | 4.63 | Oxidation reduction |
| GO:0016491 | VGI WM276 | 18 | 376 | 4.52E-04 | 3.72 | Oxidoreductase activity |
| GO:0006364 | VGII ENV152 | 34 | 13 | 1.93E-12 | 10.64 | rRNA processing |
| GO:0005730 | VGII ENV152 | 25 | 6 | 7.74E-11 | 16.95 | Nucleolus |
| GO:0003735 | VGII ENV152 | 54 | 58 | 7.37E-10 | 3.79 | Structural constituent of ribosome |
| GO:0006412 | VGII ENV152 | 71 | 117 | 4.27E-07 | 2.47 | translation |
| GO:0003723 | VGII ENV152 | 51 | 88 | 1.32E-04 | 2.36 | RNA binding |
| GO:0005506 | VGII ENV152 | 35 | 51 | 3.66E-04 | 2.79 | Iron ion binding |
| GO:0055114 | VGII ENV152 | 95 | 225 | 5.04E-04 | 1.72 | oxidation reduction |
| GO:0016627 | VGII ENV152 | 13 | 8 | 1.32E-03 | 6.61 | oxidoreductase activity, on CH-CH |
| GO:0008026 | VGII ENV152 | 24 | 32 | 2.60E-03 | 3.05 | ATP-dependent helicase activity |
| GO:0016627 | VGII R265 | 11 | 10 | 4.25E-04 | 9.6 | Oxidoreductase activity, on CH-CH |
| GO:0022900 | VGII R265 | 8 | 7 | 8.59E-03 | 9.97 | Electron transport chain |
| GO:0020037 | VGII R265 | 11 | 17 | 8.59E-03 | 5.64 | Heme binding |
| GO:0016491 | VGIII CA1873 | 33 | 361 | 9.41E-04 | 2.57 | oxidoreductase activity |
| GO:0055114 | VGIII CA1874 | 28 | 292 | 1.77E-03 | 2.69 | oxidation reduction |

Sig. depleted terms from *ex vivo* comparisons
| GO/PFAM | Isolate | count 1 | count 2 | *q* value | rel. prop | GO description |
| --- | --- | --- | --- | --- | --- | --- |
| GO:0043234 | VGII ENV152 | 20 | 194 | 1.39E-03 | 0.42 | protein complex |
| GO:0006508 | VGII ENV152 | 12 | 132 | 9.48E-03 | 0.37 | proteolysis |

### One in five *C. gattii* genes are differentially expressed between *in vitro* and *ex vivo* among the lineages

Pairwise comparisons of expression values for each lineage at different time-points (*t*0 vs *t*1, *t*3 and *t*6) identified 1,193 unique DEGs, the majority of which were upregulated *in vitro vs. ex vivo* (**Fig. 2b, Sup. Dataset 2**). About 1/3 (*n*=309) of these genes were also differentially expressed in an inter-lineage *in vitro* comparison (60% of them), leaving 215 genes that were uniquely differentially expressed *in vitro*. PCA showed large differences between *t*0 and *ex vivo* time points in both VGII isolates (**Fig. 2d**).

Genes of interest (capsule, ergosterol and laccases) were enriched for DEGs at different time points based on a Hypergeometic Test (P[X > x]) = *P*=0.007), including nine capsule biosynthesis genes (all VGII), 11 capsule attachment and cell-wall remodelling genes (all downregulated *ex vivo*), 14 ergosterol genes and both laccase genes (**Fig. 4**). Separately, many (*n*=40/97; 41%) *C. neoformans* genes that were previously found to be differentially expressed *via* microarray in the presence of amoebae and macrophages (13) were similarly modulated in *C. gattii* (111 genes with similar or different modulation between amoeba and macrophages, 97 of which had a *C. gattii* ortholog, 56 of which were also *C. gattii* DEGs between time points, and 40 had the same directionality) (**Sup. Dataset 3**).

**Figure 4.**
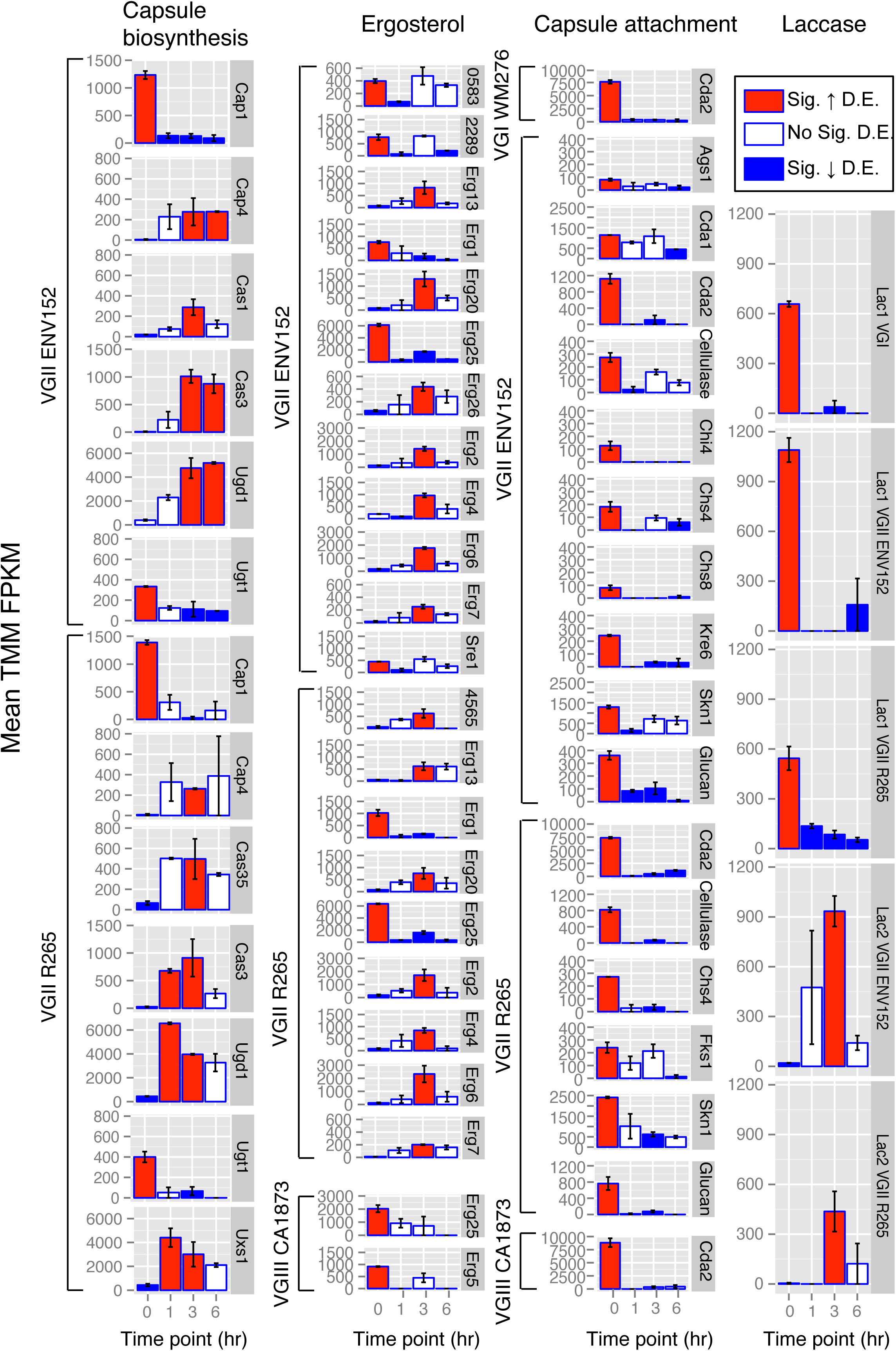
Bar charts showing mean expression (TMM FPKM) of ergosterol, capsular biosynthesis, capsular attachment and laccase genes between the five isolates of *Cryptococcus gattii* at each time point (*t*0 = *in vitro*, *t*1= 1-hour w/ BMDMs, *t*3 = 3 hours w/ BMDMs, and *t*6 = 6 hours w/ BMDMs). Red=Sig. upregulated, Blue=Sig. downregulated. Genes are considered differentially expressed where FDR *p* value<0.001 and >fourfold TMM FPKM. Error bars show the range of TMM FPKM values between the two replicates. Glucan=Glucan 1,3-β-glucosidase.

Genes differentially expressed *ex vivo* were statistically significantly enriched (two-tailed Fisher’s exact test with *q*-value FDR) for 18 GO-terms and no PFAM terms (**Table 1**). Strikingly, the oxidoreductase activity term was enriched in each lineage and isolate (apart from VGIV which had only 20 differentially expressed genes and thus no enriched terms). The ability for *C. gattii* to respond to host and environment derived oxidative stress is thus a significant feature of genes differentially expressed (both between lineages *in vitro*, and between *in vitro* and *ex vivo* conditions). Additional terms included Iron-ion binding and terms related to ribosomes (i.e. rRNA processing, structural constituent of ribosome) for VGII ENV152, perhaps indicating an increase in translational activity.

Capsule biosynthesis genes in VGII were differentially expressed in the presence of macrophages (**Fig. 4**). For example, *CAP1* and *UGT1* are downregulated by both VGII isolates, while *CAP4*, *CAS3* and *UGD1* are all upregulated at one or more of the *ex vivo* timepoints. Other capsule biosynthesis DEGs included *CAS35* (*CAS35*Δ has a decreased capsule (19)) upregulated in R265 at *t*3 *vs t*0, *UXS1* (*uxs1*Δ capsule is missing xylose (28)) upregulated in R265 at *t*1 and *t*3 *vs t*0, and *CAS1* (*cas1*Δ has a defect in capsule O-acetylation and reactivity to GXM antibodies (29)) is upregulated in VGII ENV152 at *t*3. That non-VGII lineages did not differentially express capsule synthesis genes *ex vivo* may suggest that they are less rigorously regulated (expressed in more conditions), expressed less abundantly, or are perhaps less sensitive to host-derived stresses and stimuli.

Capsule attachment and cell-wall remodelling genes were also differentially expressed *ex vivo*, predominantly by VGII isolates, and were all downregulated *ex vivo* (**Fig. 4**). For example, the chitin synthases *CHS4* and *CHS8* were upregulated *in vitro* in VGII compared with any other time points *ex vivo*. Chitin generated by such chitin synthases are converted into chitosan by the chitin deacetylases *CDA1*, *CDA2* and *CDA3*, where it constitutes an important component of the cell wall of *Cryptococcus* (30). Both *CDA1* and *CDA2* are downregulated *ex vivo* in one or more of the lineages of *C. gattii* apart from VGIV.

The four ergosterol genes CNBG0583/CNAG01003, *ERF1*, *SRE1* and *ERG25* that are upregulated in VGII compared with VGIII and VGIV *in vitro* were also upregulated in VGII *in vitro* compared with the three *ex vivo* time points, again suggesting these are downregulated during infection. However, nine additional ergosterol biosynthesis genes including *ERG2*, *ERG6*, *ERG7*, *ERG13*, *ERG20* and *ERG26* were each upregulated in the VGII isolates at three hours post infection compared with *in vitro*, suggesting these genes are activated >1 hour and <6 hours following co-incubation with macrophages.

The laccase genes that produce melanin and are upregulated in VGIV *in vitro* compared with the other lineages, are also differentially expressed in VGII between *in vitro* and *ex vivo* conditions. Specifically, *LAC1* (CNBG2144) in VGI and VGII is downregulated *ex vivo*. In contrast, *LAC2* (CNB2146) is upregulated in both VGII isolates at *t*3 compared with *t*0 – demonstrating that during infection, VGII isolates switch expression from *LAC1* to *LAC2*, perhaps owing to LAC2’s ability to remain cytoplasmic (31).

### Mouse macrophage response to *C. gattii*

Mouse BMDM expression for each of 58,716 annotated mouse transcripts (including protein-coding, pseudogenes and other non-protein coding genes) was surprisingly consistent between *in vitro* conditions and when co-incubated with any of the four lineages of *C. gattii* at any of the three time points (**Figure 5**), perhaps indicating low-rates of yeast engulfment or the general low immunogenicity of cryptococci. Only 24 upregulated DEGs and 42 downregulated DEGs were identified in total (**Sup. Dataset 4**). Of those 24 upregulated genes, five separate/unique FBJ osteosarcoma viral oncogenes (specifically *FOSB* and the truncated splice form Δ*fosb2*) belonging to the eight most highly differentially expressed genes (Log Fold Change = 3.58-7.78) were found at *t*1 for only VGII R265. *FOS* genes encode a leucine zipper protein that dimerizes with the Jun family (of which JunB is also upregulated at *t*3 in VGII ENV152) forming the AP-1 transcription factor, which in turn regulates diverse functions including cell proliferation, differentiation and transformation following the primary growth factor response. Perhaps it is therefore unsurprising that 20% of upregulated genes (*n*=11/52 redundant upregulated genes) belonged to the Early Growth Response Protein 1 family including *EGR1-201* and *EGR1-202* at *t*3 for all five *C. gattii* isolates tested, and *EGR3* at *t*1 for VGII R265 only.

**Figure 5.**
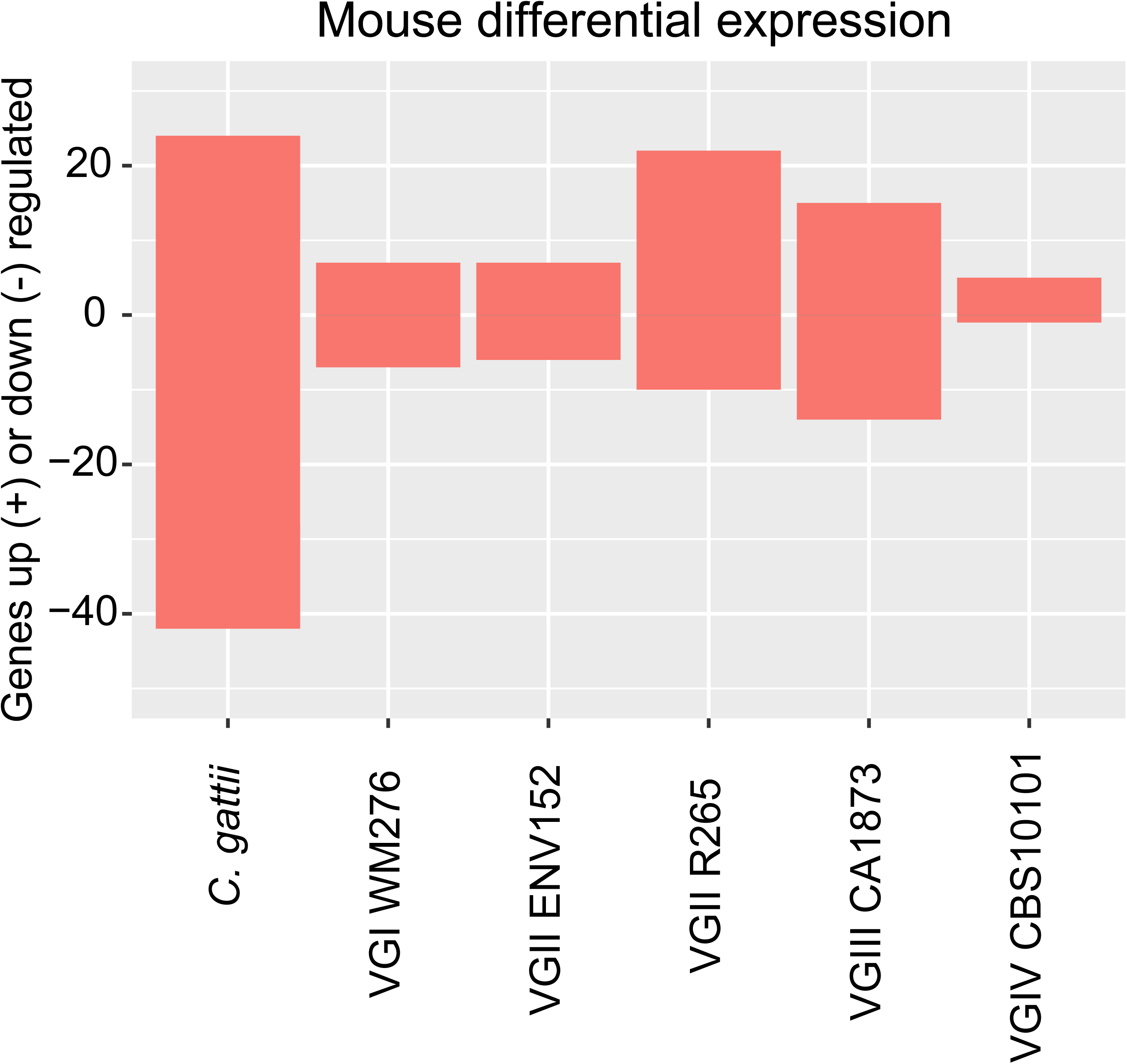
A Bar chart showing the number of Mouse genes up/downregulated by each of the five isolates of *C. gattii* compared with growth without yeast. Where the same gene is found in multiple pairwise comparisons, it is included only once in the redundant categories (only applicable for the grouped category). Positive values indicate genes that are upregulated in that isolate/lineage compared with others, while negative values (below zero) indicate genes that are downregulated in that isolate/lineage compared with others. Genes are considered differentially expressed where FDR *p* value<0.001 and >fourfold TMM FPKM.

Most of the genes downregulated in mouse macrophages during infection appear to be signalling molecules and transcription factors, including ETV5 (ETS Transcription Factor Variant 5) which is downregulated at *t*6 in all *C. gattii* isolates except VGIV. Other genes downregulated include the DENND2C proteins at *t*3 in VGII ENV152 and *t*1 in VGII R265. DENND2C proteins act in diverse intracellular signalling pathways *via* GDP/GTP exchange, with many potential downstream targets. Additionally, the DNA-binding protein ID1 (which inhibits bHLH transcription factors) was downregulated at *t*6 in VGI, VGII ENV152 and VGIII, as well as *t*1 VGII R265. Similarly, the Dual Specific Protein Phosphatase (DUSP) 6 involved in MAPK signalling was downregulated at *t*6 in VGI, VGII ENV152 and VGIII, while DUSP4 was also downregulated at *t*6 in VGI only.

## Discussion

The transcriptional responses of host and pathogen during infection can reveal key insights into their interactions and the molecular basis of pathogenicity. Transcriptional differences between distantly related isolates in culture can also reveal the impact of their genetic divergence and explain epidemiological differences. In this study, we have compared the expression profiles of five isolates belonging to the four lineages of *C. gattii* both *in vitro* between three early time points during co-incubation with mouse bone marrow derived macrophages (BMDM), and in parallel characterized the host response. Both *C. gattii* experiments suggested that lineage VGII is transcriptionally divergent compared with non-VGII lineages in terms of number of genes differentially expressed. For example, only 5% of *in vitro* DEGs belonged to non-VGII isolates, despite these isolates accounting for 30% of all pairwise comparisons made. It is likely that the loss of RNAi functionality in VGII (7, 32, 33) is partially responsible both indirectly (rewiring of gene regulatory responses in result to loss of Argonaut proteins) and perhaps even directly (mRNA not being degraded).

VGII is responsible for nearly all infections in the Pacific Northwest (PNW) (34), which is the worst recorded outbreak of *C. gattii* worldwide. VGII isolates also show a high degree (albeit across a range) of virulence traits including intracellular proliferation within macrophages (R265=1.8±0.1, ENV152=2.3±0.2), mitochondrial tubularisation percent (R265=58.4±17.6, ENV152=44.5±6.1), average phagocytosis percent (R265=31.4, ENV152=20.5), and macrophage cell death percent (R265=15.2, ENV152=12.3) (35, 36). Therefore, isolates within lineages demonstrate high phenotypic variability. Although the sequenced isolates in this study are representative of each linage, comparing multiple isolates from each lineage would help delineate the intra *vs.* inter lineage variation in gene expression.

The capsule biosynthesis pathway in *C. gattii* constitutes a complex trait controlled by numerous genes. Many of these genes presented varying degrees of expression among the four lineages, in particular VGII, suggesting diverse mechanisms to maintain and perhaps even diversify the properties of the capsule. For example, *CAS3* is upregulated by VGII isolates at various time-points *ex vivo* compared with *in vitro*. Gene expression for capsule genes was variable both in benign (rich media) and stressful (co-incubation with BMDM) environments. *CAS3* has previously been identified as upregulated in VGII R265 compared with low-virulent VGII isolate R272 under conditions of carbon and nitrogen starvation (16). However, we found no evidence of differential expression of *CAS3* between VGII R265 and VGII ENV152, or indeed any other isolate – possibly indicating a uniqueness of R272 expression, or experimental differences between studies. Separately, the capsule biosynthesis gene *CAP2* is upregulated by VGIII, while *CAP1* is downregulated by VGI and VGIII, suggesting a lineage transition from expressing one gene to another. Capsule attachment DEGs (in comparison to capsule biosynthesis genes) were all downregulated *ex vivo*, suggesting these do not play an active role during infection.

Ergosterol in the membrane of *C. gattii* is a key target for numerous anti-fungal drugs including fluconazole and amphotericin B (20). VGII presented higher expression for genes involved in ergosterol production *in vitro*, including an upregulation of *ERG1*, *ERG25*, *SRE1* and an NADPH-ferrihemoprotein reductase. Mutants for each of these genes show a range of defects in the presence of antifungals or hypoxia(21–24). *ERG1*, *ERG25* and *SRE1* were also upregulated in VGII *in vitro* compared with the three *ex vivo* time points, suggesting these genes can be switched off during non-drug related stresses. However, a further nine ergosterol genes were upregulated between 1 and 6 hours post infection, including *ERG2*, *ERG6*, *ERG7*, *ERG13*, *ERG20* and *ERG26*, suggesting that this pathway is active during infection.

Laccase production (*LAC1* and *LAC2*) in both *C. gattii* and *C. neoformans* is controlled by two cell wall enzymes that possesses a broad spectrum of activity oxidizing both polyphenolic compounds and iron (25). VGIV upregulates both genes compared with other lineages *in vitro*. *LAC1* is downregulated in VGI and VGII *ex vivo* compared with *in vitro*, and instead *LAC2* is upregulated in both VGII isolates at *t*3 compared with *t*0 – suggesting that *C. gattii* VGII switches expression from one laccase gene to the other during infection, perhaps due to changing concentrations or requirements for metabolism of lactose and galactose, or production of melanin. In *C. neoformans*, *LAC1* and *LAC2* (along with capsule genes) are part of the Gpa1-cAMP pathway, which regulates capsule and melanin production using L-dopa as a substrate (37). We hypothesise that VGII uses *LAC2* to regulate growth and glucose responses (and potentially virulence) instead of *LAC1*. In *C. neoformans*, *LAC1* is localised to the cell wall, whereas *LAC2* is cytoplasmic, but is capable of localising to the cell wall (31); therefore, *C. gattii LAC2* could behave similarly to its *C. neoformans* counterpart, which may have increased versatility during infection of macrophages.

We also identified differential expression in VGII lineage-specific genes such as an MFS transporter and an oxidoreductase that may have contributed to the enrichment of ‘oxidative reduction’, which was identified in both *in vitro* comparisons and *in vitro vs ex vivo* comparisons. This term includes a wide-range of functions, pathways, and genes including ferric reductases, metalloreductases, nitric oxide dioxygenases, acidic laccases, oxidoreductases and various dehydrogenases. Previously, an upregulation of genes involved in oxidative stress has been identified in *C. neoformans* isolate H99 co-incubated with J774A macrophage-like cell line for 16 hours *vs in vitro* conditions (12), and in *C. gattii* VGII isolates R265 vs R272 in carbon and nitrogen starvation (16).The ability for *C. gattii* to respond to host and environment derived oxidative stress is well described – and it is consistent that genes involved in these processes should also be enriched between isolates and lineages at the expression level *in vitro* and among lineages at different time points *ex vivo*.

Finally, we found that mouse macrophages respond to *C. gattii* by upregulating FosB/Jun/Egr1 regulatory proteins at early time points, which may trigger differentiation and cell division of macrophages. We found little evidence for differential expression induced by different *C. gattii* lineages suggesting that the macrophage response to *C. gattii* is the same across lineages. Our study highlights the breadth of expression profiles amongst the lineages of *C. gattii*, and the diversity of transcriptional responses at this host-pathogen interface, some of which may be the cause for differences in phenotypic and clinical manifestations noted between lineages (38).

## Acknowledgments

RAF was supported by a Wellcome Trust-Massachusetts Institute of Technology (MIT) Postdoctoral Fellowship. MCF and JR were supported by a Medical Research Grant MR/K000373/1. All RNA-seq data for Mouse macrophages and *Cryptococcus gattii* has been deposited in the Short Read Achieve under the accession SRP128463. RCM is supported by a Wolfson Royal Society Research Merit Award and by funding from the European Research Council under the European Union’s Seventh Framework Programme (FP/2007-2013)/ERC Grant Agreement No. 614562

## Figure legends

**Figure S1. Alignments of *C. gattii* and Mouse transcripts to the Core Eukaryotic Genes (CEGs) suggesting mostly complete *C. gattii* gene-sets and high-quality of Mouse gene-set mm10 p4.** Only VGIIa R265 and Mouse mm10p4 were used in this study.

**Figure S2. RNA was extracted from all four lineages of *C. gattii* in mouse macrophages at 1hr, 3hr and 6hr post infection, as well *C. gattii* and macrophages *in vitro* (*t*0).** The percent of sequenced reads deriving from the mouse macrophage (above) and *C. gattii* (below).

**Figure S3. Heat maps of all differentially expressed genes (FDR *p* value<0.001 and >fourfold change of trimmed mean of M-values (TMM) normalized Fragments Per Kilobase of transcript per Million mapped reads (FPKM)) of *C. gattii* transcripts *in vitro* (*t*0) and during infection of mouse macrophages at 1hr, 3hr and 6hr.**

**Figure S4. Subsets (75%, 50% and 25%) of *C. gattii* data were used to recall differential expression and compare with the full dataset. X-axis categories for each Bar chart are each of the five isolates (combining *t*1, *t*3, *t*6), and *in vitro* (combining each isolate at *t*0). (a)** The total number of differentially expressed genes changes (either going up or down) for isolates *ex vivo*, but not *in vitro*. **(b)** The percent of genes found in the full dataset (i.e. a proxy for true positives). >75% for all categories using 75% subsets, >50% for all categories using 50% subsets, and >25% for all categories using 25% subsets. **(c)** The number of genes found only in the full dataset (i.e. a proxy for false negatives). VGI, VGIII and VGIV re-identified most of the genes also found with their full datasets, while VGII and *in vitro* conditions recovered fewer genes found in the full dataset as the subset becomes smaller. **(d)** The number of genes not found the full dataset (i.e. a proxy for false positives). Previously unidentified genes either increased or decreased as subset size decreased, with VGIV being the most robust and VGIII the least.

**Figure S5. A paralogous cluster of Cas3 and Cas31 from a previously described study of 16 isolates** (7) **had their sequences aligned using MUSCLE v3.8.31** (40)**, and a neighbor-joining tree constructed using PAUP version 4.0b10** (41) **to decipher orthologs.** *Cn*=*Cryptococcus neoformans*, *Cg*=*Cryptococcus gattii*. Note that VGIV has no Cas31 gene. Expression levels (TMM FPKM) for five isolates are shown below the dendrogram.

**Table S1. (Tab 1) Contamination from *E. coli* K-12 MG1655, Suicide vector pCD-RAsl1 and Cloning vector pMJ016c identified by BLASTn searches to the NR database. (Tab2) Reads aligning to either mouse GRCm38 p4 mm10 gene sets or genome, or R265 updated gene set or genome.** ARD=Average Read Depth across genes.

**Supplemental Datasets. Differentially expressed genes. (Tab 1-4)** All differentially expressed genes between five *C. gattii* isolates *in vitro*. **(Tab 1)** Details of every gene differentially expressed. **(Tab 2)** Gene annotation, position in genome, and associated PFAM. Orthogroup number refers to ortholog groups from 15 *C. gattii* and *C. neoformans* H99 genomes comparisons (7). **(Tab 3)** Gene ontology terms assigned to differentially expressed genes. **(Tab 4)** Differentially expressed genes were found in multiple pairwise comparisons. (**Tab 5-8**) All differentially expressed genes by each of the five *C. gattii* isolates at different time points *(t*0*, t*3*, t*6) with co-incubation with mouse macrophages. **(Tab 5)** Details of every gene differentially expressed. **(Tab 6)** Gene annotation, position in genome, and associated PFAM. Orthogroup number refers to ortholog groups from 15 *C. gattii* and *C. neoformans* H99 genomes comparisons (7). **(Tab 7)** Gene ontology terms assigned to differentially expressed genes. **(Tab 8)** Differentially expressed genes were found in multiple pairwise comparisons. (**Tab 9-10**) Expression values and overlap with previously identified differentially expressed genes in *Acanthamoeba castellanii* and Macrophages (13). **(Tab 9)** *C. neoformans* genes with similar modulation patterns after interaction of the fungus with amoebae and murine macrophages. (**Tab 10**) *C. neoformans* genes with different modulation patterns after interaction of the fungus with amoeba and murine macrophages. (**Tab 11**) Genes differentially expressed by Mouse macrophages. LogFc=Log Fold Change. LogCPM=Log Counts Per Million. C1=VGIV CBS10101, C2=VGII R265, C3=VGII ENV152, C4=CA1873, C5=VGI WM276. A and B = Replicates.

## Methods

### Macrophage and *Cryptococcus gattii* infection assay

*C. gattii* was grown in accordance with previous work(26). Briefly, five strains of *C. gattii* were grown in RPMI/C10 w/HI fetal bovine serum (FBS) media, and pre-incubated at 37°C in C10 media for 1.5hrs before infection. Bone marrow derived macrophages (BMDM) were derived from bone marrow cells collected from the femur and tibia of C57BL/6 female mice. All mouse work was performed in accordance with the Institutional Animal Care and Use Committees (IACUC) and with relevant guidelines at the Broad Institute and Massachusetts Institute of Technology, with protocol 0615-058-1. Primary bone marrow cells were grown in “C10” media as previously described^42^ and supplemented with macrophage colony stimulating factor (M-CSF) (ThermoFisher Scientific) at final concentration of 10 ng/ml to promote differentiation into macrophages.

For the infection experiment, dilutions of 7.5E+5 cells were made for BMDM and *C. gattii* strains. BMDM were spun at 500rcf at 37°C for 2 minutes to adhere to plates, and inoculated with *C. gattii* strains using a multiplicity of infection (MOI) of 1 macrophage to 2 *C. gattii*. Macrophages with *C. gattii* cells were centrifuged at 500rcf at 37°C for 2 minutes. Next, cells were incubated at 37°C with 5% CO_2_ for 1hr, 3hr and 6hr post centrifuge. In addition, we grew *C. gattii in vitro*, and BMDM without *C. gattii*, in duplicate with the same C10 media. At the end of the time points, the media was removed, a lysis buffer added, and placed in a −70°C freezer for RNA extraction.

RNA was extracted from population samples using the Qiagen RNeasy mini kit. All samples were subjected to 3 minutes of bead beating with .5mm zirconia glass beads (BioSpec Products) in a bead mill. Libraries were generated with the TruSeq Stranded mRNA Library Prep kit (Illumina).

### RNA-seq and data analysis

All samples were sequenced on Illumina HiSeq 2500 to generate strand-specific paired-end reads of 38nt length. Raw reads from each sample were sequenced on multiple lanes, so were merged into individual replicates. Using Blast searches to the nr database, we identified the following contaminants in each of the samples: *E. coli* K-12 MG1655, Suicide vector pCD-RAsl1 and Cloning vector pMJ016c (**Table S1**). These contaminants were excluded from all of the read sets based on Bowtie2 (43) alignments. We next aligned all reads to mouse GRCm38 p4 mm10 transcripts sets (including protein coding, pseudogenes, and other categories of non-protein coding genes) and genome (44) using Bowtie2 (43). We took all reads that aligned to neither the mouse genome or gene sets, and aligned to the R265 updated gene set (7) as well as the R265 genome (**Table S2**). Both VGIIa R265 and mm10p4 gene-sets were found to be largely complete in terms of Core Eukaryotic Gene coverage (**Fig. S1**).

Reads that aligned only to *C. gattii*, or only to mm10, had their transcript abundance estimated using RSEM (45) and differential expression (FDR *p*-value < 0.001 and > 4 fold change of TMM normalized FPKM) using EdgeR (17) through the Trinity version r20140413p1 (46) pipeline. Specifically, the pipeline uses the EdgeR quantile-adjusted conditional maximum likelihood (qCML) method after estimating common dispersion and tagwise dispersions using the Cox-Reid profile-adjusted likelihood method (also implemented in EdgeR). For *Cryptococcus* strains, we estimated transcript abundance and differential expression in the same way after 1) merged time-point (i.e. calculations on each isolate only), and 2) merged isolates (i.e. calculations on each time point only). Merging by isolates had far weaker correlation between ‘replicates’ than merging time points, indicating *in vivo* conditions were more influential on overall expression values than lineage/strain specific differences. Principle Component Analysis (PCA) was performed using SmartPCA from EIGENSOFT v4.0 (47), where each gene was given the value 0 for non-DEG, and the value 1 for DEG (upregulated).

Orthologs to genes of known function in *C. neoformans* H99 were identified in *C. gattii* R265 using OrthoMCL (48) across 16 isolates as previously described (7). Genes of interest (capsule biosynthesis, capsule attachment, and ergosterol) that fell within paralogous clusters had the sequences of 16 isolates described previously (7) aligned using MUSCLE v3.8.31 (40), and a neighbor-joining tree constructed using PAUP version 4.0b10 (41) to decipher orthologs. Synteny was visualised using Synima (39).

Of the 58,716 annotated mouse transcripts, 39,229 (67%) had evidence of expression in one or more samples (TMM FPKM > 1). Prior to differential expression analysis, we excluded 3,596 mouse transcripts that any *C. gattii in vitro* RNA (i.e. no macrophage present) aligned to. Surprisingly, some mouse transcripts were very highly expressed in *C. gattii* only samples (e.g. glutamate receptor interacting protein 2 (Grip2-205) had TMM FPKM > 15 thousand for every *C. gattii* isolate, and only between 10 and 874 for *t*1, *t*3 and *t*6, when mouse macrophages were actually present), perhaps owing to sequence similarity of some 30mers, or inaccuracies in the mouse gene-set. Applying the same threshold of FDR *p*-value < 0.001 and > 4 fold change of TMM normalized FPKM) using EdgeR (17) identified 24 genes that were upregulated during infection by 1 or more isolate of *C. gattii*, and 42 genes that were downregulated during infection by 1 or more isolate of *C. gattii*.

Enrichment analyses for PFAM and GO terms previously assigned (7) were conducted using two-tailed Fisher’s exact test with *q*-value FDR. Multiple testing corrections were performed with the Storey-Tibshirani (49) method (requiring *q*-value <0.05). For enrichment tests, we excluded PFAM and GO terms related to transposable elements and domains of unknown function.

### Impact of read depth on differential expression

Based on data quantity / transcriptome coverage alone, the host-response and *in vitro C. gattii* lineage expression differences should be most robust, while *C. gattii ex vivo* expression changes should be less robust. To explore the coverage and sensitivity of our *C. gattii* RNAseq data and pipeline, we first made subsets (75%, 50% and 25%) of our *C. gattii* data and re-called differential expression (**Fig. S3**). The number of genes differentially expressed between isolates *in vitro* conditions remained most consistent in terms of number of differentially expressed genes following sub-setting (100% = 1,208, 75%=1,205, 50%=1,147, 25%=800) (**Fig. S3A**).

For all *C. gattii* datasets (those from *in vitro* conditions, and individual isolates between conditions), the number of genes re-identified in the 75% subset was between 79%-100% (VGIV CBS10101 had the same 20 genes in both Dataset sizes) (**Fig. S3B**). Conversely, the number of genes identified only in the full datasets increased in the remaining subsets, with the most pronounced being VGIIa ENV152 losing the identification of 476 genes in the 75% subset, possibly indicating a lack of data for this isolate (**Fig. S3C**). We also assessed each subset for genes that were not found in the full set (i.e. were unique to the subsets, **Fig. S3D**). VGII, VGIV and the *in vitro* datasets had fewer unique genes as subsets became smaller, while VGI and VGIII had an increase as subsets became smaller.

For *in vitro* conditions, between 10%-15% of genes were unique in their subsets, while for VGII ENV152, it was between 1%-5%. For isolates with very few identified differentially expressed genes (such as VGIV CBS10101 and VGIII CA1873), the numbers of unique genes approached or even exceeded the number found in the full dataset. Given this variation, it is therefore likely that most of the *ex vivo C. gattii* datasets would benefit from deeper RNAseq data, and that the *in vitro* expression values are more robust than *ex vivo* data. Nevertheless, the majority of differential expressed genes were consistent between the larger subsets of data

## References

1. Fisher MC, Henk DA, Briggs CJ, Brownstein JS, Madoff LC, McCraw SL, Gurr SJ. 2012. Emerging fungal threats to animal, plant and ecosystem health. Nature 484:186–194.

2. Farrer RA, Fisher MC. 2017. Describing genomic and epigenomic traits underpinning emerging fungal pathogens. Adv Genet 100:73–140.

3. Farrer RA, Weinert LA, Bielby J, Garner TWJ, Balloux F, Clare F, Bosch J, Cunningham AA, Weldon C, Preez LH du, Anderson L, Pond SLK, Shahar-Golan R, Henk DA, Fisher MC. 2011. Multiple emergences of genetically diverse amphibian-infecting chytrids include a globalized hypervirulent recombinant lineage. Proc Natl Acad Sci 108:18732–18736.

4. Farrer RA, Henk DA, Garner TWJ, Balloux F, Woodhams DC, Fisher MC. 2013. Chromosomal copy number variation, selection and uneven rates of recombination reveal cryptic genome diversity linked to pathogenicity. PLoS Genet 9:e1003703.

5. Springer DJ, Phadke S, Billmyre B, Heitman J. 2012. *Cryptococcus gattii*, no longer an accidental pathogen? Curr Fungal Infect Rep 6:245–256.

6. Fraser JA, Giles SS, Wenink EC, Geunes-Boyer SG, Wright JR, Diezmann S, Allen A, Stajich JE, Dietrich FS, Perfect JR, Heitman J. 2005. Same-sex mating and the origin of the Vancouver Island Cryptococcus gattii outbreak. Nature 437:1360–1364.

7. Farrer RA, Desjardins CA, Sakthikumar S, Gujja S, Saif S, Zeng Q, Chen Y, Voelz K, Heitman J, May RC, Fisher MC, Cuomo CA. 2015. Genome Evolution and Innovation across the Four Major Lineages of Cryptococcus gattii. mBio 6.

8. Li H, Johnson AD. 2010. Evolution of Transcription Networks — Lessons from Yeasts. Curr Biol 20:R746–R753.

9. Fraser HB, Levy S, Chavan A, Shah HB, Perez JC, Zhou Y, Siegal ML, Sinha H. 2012. Polygenic cis-regulatory adaptation in the evolution of yeast pathogenicity. Genome Res 22:1930–1939.

10. Cooke DEL, Cano LM, Raffaele S, Bain RA, Cooke LR, Etherington GJ, Deahl KL, Farrer RA, Gilroy EM, Goss EM, Grünwald NJ, Hein I, MacLean D, McNicol JW, Randall E, Oliva RF, Pel MA, Shaw DS, Squires JN, Taylor MC, Vleeshouwers VGAA, Birch PRJ, Lees AK, Kamoun S. 2012. Genome analyses of an aggressive and invasive lineage of the Irish potato famine pathogen. PLoS Pathog 8:e1002940.

11. Ma H, Hagen F, Stekel DJ, Johnston SA, Sionov E, Falk R, Polacheck I, Boekhout T, May RC. 2009. The fatal fungal outbreak on Vancouver Island is characterized by enhanced intracellular parasitism driven by mitochondrial regulation. Proc Natl Acad Sci U S A 106:12980–12985.

12. Fan W, Kraus PR, Boily M-J, Heitman J. 2005. *Cryptococcus neoformans* gene expression during murine macrophage infection. Eukaryot Cell 4:1420–1433.

13. da Derengowski L S, Paes HC, Albuquerque P, Tavares AHFP, Fernandes L, Silva-Pereira I, Casadevall A. 2013. The transcriptional response of *Cryptococcus neoformans* to ingestion by *Acanthamoeba castellanii* and macrophages provides insights into the evolutionary adaptation to the mammalian host. Eukaryot Cell 12:761–774.

14. Janbon G, Ormerod KL, Paulet D, Iii EJB, Yadav V, Chatterjee G, Mullapudi N, Hon C-C, Billmyre RB, Brunel F, Bahn Y-S, Chen W, Chen Y, Chow EWL, Coppée J-Y, Floyd-Averette A, Gaillardin C, Gerik KJ, Goldberg J, Gonzalez-Hilarion S, Gujja S, Hamlin JL, Hsueh Y-P, Ianiri G, Jones S, Kodira CD, Kozubowski L, Lam W, Marra M, Mesner LD, Mieczkowski PA, Moyrand F, Nielsen K, Proux C, Rossignol T, Schein JE, Sun S, Wollschlaeger C, Wood IA, Zeng Q, Neuvéglise C, Newlon CS, Perfect JR, Lodge JK, Idnurm A, Stajich JE, Kronstad JW, Sanyal K, Heitman J, Fraser JA, Cuomo CA, Dietrich FS. 2014. Analysis of the genome and transcriptome of *Cryptococcus neoformans var. grubii* reveals complex RNA expression and microevolution leading to virulence attenuation. PLOS Genet 10:e1004261.

15. Chen Y, Toffaletti DL, Tenor JL, Litvintseva AP, Fang C, Mitchell TG, McDonald TR, Nielsen K, Boulware DR, Bicanic T, Perfect JR. 2014. The *Cryptococcus neoformans* transcriptome at the site of human meningitis. mBio 5.

16. Ngamskulrungroj P, Price J, Sorrell T, Perfect JR, Meyer W. 2011. *Cryptococcus gattii* virulence composite: candidate genes revealed by microarray analysis of high and less virulent Vancouver island outbreak strains. PloS One 6:e16076.

17. Robinson MD, McCarthy DJ, Smyth GK. 2010. EdgeR: a Bioconductor package for differential expression analysis of digital gene expression data. Bioinformatics 26:139–140.

18. O’Meara TR, Alspaugh JA. 2012. The *Cryptococcus neoformans* capsule: a sword and a shield. Clin Microbiol Rev 25:387–408.

19. Moyrand F, Chang YC, Himmelreich U, Kwon-Chung KJ, Janbon G. 2004. Cas3p belongs to a seven-member family of capsule structure designer proteins. Eukaryot Cell 3:1513–1524.

20. Santos JRA, Gouveia LF, Taylor ELS, Resende-Stoianoff MA, Pianetti GA, César IC, Santos DA. 2012. Dynamic interaction between fluconazole and amphotericin B against *Cryptococcus gattii*. Antimicrob Agents Chemother 56:2553–2558.

21. Tsai H-F, Bard M, Izumikawa K, Krol AA, Sturm AM, Culbertson NT, Pierson CA, Bennett JE. 2004. Candida glabrata erg1 Mutant with Increased Sensitivity to Azoles and to Low Oxygen Tension. Antimicrob Agents Chemother 48:2483–2489.

22. Kim J, Cho Y-J, Do E, Choi J, Hu G, Cadieux B, Chun J, Lee Y, Kronstad JW, Jung WH. 2012. A defect in iron uptake enhances the susceptibility of *Cryptococcus neoformans* to azole antifungal drugs. Fungal Genet Biol FG B 49:955–966.

23. Blosser SJ, Merriman B, Grahl N, Chung D, Cramer RA. 2014. Two C4-sterol methyl oxidases (Erg25) catalyse ergosterol intermediate demethylation and impact environmental stress adaptation in Aspergillus fumigatus. Microbiol Read Engl 160:2492–2506.

24. Chun CD, Liu OW, Madhani HD. 2007. A link between virulence and homeostatic responses to hypoxia during infection by the human fungal pathogen Cryptococcus neoformans. PLoS Pathog 3:e22.

25. Zhu X, Williamson PR. 2004. Role of laccase in the biology and virulence of Cryptococcus neoformans. FEMS Yeast Res 5:1–10.

26. Voelz K, Johnston SA, Smith LM, Hall RA, Idnurm A, May RC. 2014. ‘Division of labour’ in response to host oxidative burst drives a fatal *Cryptococcus gattii* outbreak. Nat Commun 5:5194.

27. Wang Y, Casadevall A. 1994. Susceptibility of melanized and nonmelanized *Cryptococcus neoformans* to nitrogen-and oxygen-derived oxidants. Infect Immun 62:3004–3007.

28. Kozel TR, Levitz SM, Dromer F, Gates MA, Thorkildson P, Janbon G. 2003. Antigenic and biological characteristics of mutant strains of *Cryptococcus neoformans* lacking capsular O acetylation or xylosyl side chains. Infect Immun 71:2868–2875.

29. Moyrand F, Klaproth B, Himmelreich U, Dromer F, Janbon G. 2002. Isolation and characterization of capsule structure mutant strains of Cryptococcus neoformans. Mol Microbiol 45:837–849.

30. Baker LG, Specht CA, Donlin MJ, Lodge JK. 2007. Chitosan, the deacetylated form of chitin, is necessary for cell wall integrity in *Cryptococcus neoformans*. Eukaryot Cell 6:855–867.

31. Missall TA, Moran JM, Corbett JA, Lodge JK. 2005. Distinct Stress Responses of Two Functional Laccases in Cryptococcus neoformans Are Revealed in the Absence of the Thiol-Specific Antioxidant Tsa1. Eukaryot Cell 4:202–208.

32. D’Souza CA, Kronstad JW, Taylor G, Warren R, Yuen M, Hu G, Jung WH, Sham A, Kidd SE, Tangen K, Lee N, Zeilmaker T, Sawkins J, McVicker G, Shah S, Gnerre S, Griggs A, Zeng Q, Bartlett K, Li W, Wang X, Heitman J, Stajich JE, Fraser JA, Meyer W, Carter D, Schein J, Krzywinski M, Kwon-Chung KJ, Varma A, Wang J, Brunham R, Fyfe M, Ouellette BFF, Siddiqui A, Marra M, Jones S, Holt R, Birren BW, Galagan JE, Cuomo CA. 2011. Genome variation in *Cryptococcus gattii*, an emerging pathogen of immunocompetent hosts. mBio 2:e00342–00310.

33. Wang X, Hsueh Y-P, Li W, Floyd A, Skalsky R, Heitman J. 2010. Sex-induced silencing defends the genome of *Cryptococcus neoformans via* RNAi. Genes Dev 24:2566–2582.

34. Byrnes EJ, Li W, Lewit Y, Ma H, Voelz K, Ren P, Carter DA, Chaturvedi V, Bildfell RJ, May RC, Heitman J. 2010. Emergence and pathogenicity of highly virulent *Cryptococcus gattii* genotypes in the northwest United States. PLoS Pathog 6:e1000850.

35. Farrer RA, Voelz K, Henk DA, Johnston SA, Fisher MC, May RC, Cuomo CA. 2016. Microevolutionary traits and comparative population genomics of the emerging pathogenic fungus *Cryptococcus gattii*. Phil Trans R Soc B 371:20160021.

36. Voelz K, Ma H, Phadke S, Byrnes EJ, Zhu P, Mueller O, Farrer RA, Henk DA, Lewit Y, Hsueh Y-P, Fisher MC, Idnurm A, Heitman J, May RC. 2013. Transmission of hypervirulence traits via sexual reproduction within and between lineages of the human fungal pathogen *Cryptococcus gattii*. PLoS Genet 9.

37. Pukkila-Worley R, Gerrald QD, Kraus PR, Boily M-J, Davis MJ, Giles SS, Cox GM, Heitman J, Alspaugh JA. 2005. Transcriptional network of multiple capsule and melanin genes governed by the *Cryptococcus neoformans* cyclic AMP cascade. Eukaryot Cell 4:190–201.

38. Chen SC-A, Slavin MA, Heath CH, Playford EG, Byth K, Marriott D, Kidd SE, Bak N, Currie B, Hajkowicz K, Korman TM, McBride WJH, Meyer W, Murray R, Sorrell TC, Australia and New Zealand Mycoses Interest Group (ANZMIG)-Cryptococcus Study. 2012. Clinical manifestations of *Cryptococcus gattii* infection: determinants of neurological sequelae and death. Clin Infect Dis 55:789–798.

39. Farrer RA. 2017. Synima: a Synteny imaging tool for annotated genome assemblies. BMC Bioinformatics 18:507.

40. Edgar RC. 2004. MUSCLE: a multiple sequence alignment method with reduced time and space complexity. BMC Bioinformatics 5:113.

41. Yang Z. 2007. PAML 4: phylogenetic analysis by maximum likelihood. Mol Biol Evol 24:1586–1591.

42. Stubbs MC, Kim YM, Krivtsov AV, Wright RD, Feng Z, Agarwal J, Kung AL, Armstrong SA. 2008. MLL-AF9 and FLT3 cooperation in acute myelogenous leukemia: development of a model for rapid therapeutic assessment. Leukemia 22:66–77.

43. Langmead B, Salzberg SL. 2012. Fast gapped-read alignment with Bowtie 2. Nat Methods 9:357–359.

44. Blake JA, Bult CJ, Eppig JT, Kadin JA, Richardson JE, Group TMGD. 2014. The Mouse Genome Database: integration of and access to knowledge about the laboratory mouse. Nucleic Acids Res 42:D810–D817.

45. Li B, Dewey CN. 2011. RSEM: accurate transcript quantification from RNA-Seq data with or without a reference genome. BMC Bioinformatics 12:323.

46. Grabherr MG, Haas BJ, Yassour M, Levin JZ, Thompson DA, Amit I, Adiconis X, Fan L, Raychowdhury R, Zeng Q, Chen Z, Mauceli E, Hacohen N, Gnirke A, Rhind N, di Palma F, Birren BW, Nusbaum C, Lindblad-Toh K, Friedman N, Regev A. 2011. Full-length transcriptome assembly from RNA-Seq data without a reference genome. Nat Biotech 29:644–652.

47. Patterson N, Price AL, Reich D. 2006. Population Structure and Eigenanalysis. PLOS Genet 2:e190.

48. Li L, Stoeckert CJ, Roos DS. 2003. OrthoMCL: identification of ortholog groups for eukaryotic genomes. Genome Res 13:2178–2189.

49. Storey JD, Tibshirani R. 2003. Statistical significance for genomewide studies. Proc Natl Acad Sci U S A 100:9440–9445.

